# An Extension of the Fri Framework for Calcium Transient Detection

**DOI:** 10.1101/029751

**Authors:** Stephanie Reynolds, Caroline S. Copeland, Simon R. Schultz, Pier Luigi Dragotti

## Abstract

Two-photon calcium imaging of the brain allows the spatiotemporal activity of neuronal networks to be monitored at cellular resolution. In order to analyse this activity it must first be possible to detect, with high temporal resolution, spikes from the time series corresponding to single neurons. Previous work has shown that finite rate of innovation (FRI) theory can be used to reconstruct spike trains from noisy calcium imaging data. In this paper we extend the FRI framework for spike detection from calcium imaging data to encompass data generated by a larger class of calcium indicators, including the genetically encoded indicator GCaMP6s. Furthermore, we implement least squares model-order estimation and perform a noise reduction procedure (‘pre-whitening’) in order to increase the robustness of the algorithm. We demonstrate high spike detection performance on real data generated by GCaMP6s, detecting 90% of electrophysiologically-validated spikes.

***Index Terms***— Calcium imaging, Calcium transient detection, Finite rate of innovation, GCaMP6s

## 1. INTRODUCTION

Optical imaging of populations of neurons at cellular resolution may prove crucial to developing our understanding of the function of the brain. In order to analyse the spatiotemporal activity of neuronal networks, one must first be able to detect, with high temporal precision, the time at which action potentials (or spikes) were fired from individual neurons.

As the concentration of intracellular free calcium is a reliable indicator of spiking activity, several optical imaging methods rely upon calcium-sensitive fluorescent indicators (calcium indicators) which visualise spiking activity via a change in their fluorescence intensity. Recent advances in protein engineering have produced genetically encoded calcium indicators (GECIs) which have, for the first time, exceeded the sensitivity of the traditionally used synthetic indicators [1]. GECIs have a proven capability for imaging the same *in vivo* neuronal populations over multiple weeks [2] and they can be targeted to selected cellular and subcellular compartments. Due to the above advantages of GECIs, they are becoming the preferred tool for calcium imaging experiments. Spike detection algorithms which are suited to the kinetics of these indicators are thus required.

A spike in a neuron produces a pulse with a characteristic shape in that neuron’s fluorescence signal, this pulse is referred to as a calcium transient. Several algorithms employ template-matching approaches which locate portions of the fluorescence signal which correspond to the expected pulse template [3, 4, 5]. The performance of such algorithms which include strict assumptions on the template’s amplitude [5] deteriorates on real data, in which transient amplitudes vary greatly. Vogelstein et al. developed a fast algorithm that performs a *maximum a posteriori* estimation to infer the most likely spike train given the imaging data and a model of intracellular calcium dynamics [6].

In [7], Oñativia et al. exploited the fact that calcium imaging data, which can be modelled as streams of decaying exponentials, are a class of signals with a finite rate of innovation (FRI) [8]. They used FRI theory to develop a fast spike detection algorithm which demonstrated both high accuracy and high temporal precision when detecting spikes from calcium imaging data generated by the synthetic dye Oregon Green BAPTA 1-AM (OGB-1).

The main focus of this paper is to extend the FRI framework for calcium transient detection to be used on a larger class of calcium indicators. Oñativia et al. modelled the characteristic pulse template as an instantaneous rise and an exponential decay. This is a good approximation for some calcium indicators, but not for those with a slow rise such as the GECI GCaMP6s, which has already been shown to be highly useful for the study of neuronal networks [9, 1, 10]. Transients generated by GCaMP6s take 200-300ms to reach peak amplitude and thus will be detected with low temporal precision by algorithms which assume an instantaneous rise time.

In Section 2 we generalise the FRI framework for calcium transient detection for a pulse template which approximates the dynamics of a larger class of calcium indicators. In Section 2.2 we introduce a method to increase the robustness of spike detection from noisy fluorescence signals and in Section 2.3 we outline our least squares model-order estimation framework. Finally, in Section 3 we demonstrate the performance of the modified FRI algorithm on real data generated by the calcium indicator GCaMP6s.

## 2. FINITE RATE OF INNOVATION THEORY APPLIED TO CALCIUM TRANSIENT DETECTION

A spike in a neuron produces a calcium transient with a characteristic pulse shape in the corresponding neuron’s fluorescence signal. This signal can therefore be modelled as a convolution of the spike train 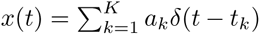 with the known characteristic pulse shape *p*(*t*), such that

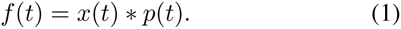

Finite Rate of Innovation (FRI) theory is a framework for the sampling and reconstruction of signals that can be completely defined by a finite number of free parameters. The signal *f*(*t*) is an example of such a signal as it is completely defined by the parameter set 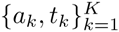.

In [7], Oñativia et al. develop an FRI algorithm to detect the time points of calcium transients whose shape is characterised by an instantaneous rise and an exponential decay. They initially filter the signal *f*(*t*) with an exponential reproducing kernel and compute weighted finite differences of the samples. These operations allow the authors to transform the problem from one of estimating the time points of calcium transients to the classical FRI problem of retrieving the locations of a stream of Diracs. Transforming the problem into a classical one in the FRI framework enables the authors to use FRI methods (see [11]) to retrieve the time points of the calcium transients.

We now focus on extending the FRI framework for calcium transient detection to those whose shape is characterised by a slower (not instantaneous) rise and an exponential decay. We model their characteristic pulse shape as

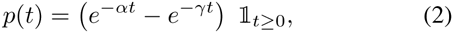
where *α* and γ are known parameters. In order to use the FRI framework for this pulse shape, we first need to identify the filtering scheme that transforms the estimation problem into sampling and reconstructing a stream of Diracs.

### Proposition 1.

*Filtering f(t) = x(t) * p(t) with the scheme in (3)*

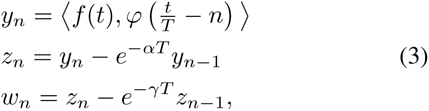

*is analogous to sampling the stream of Diracs x(t) with the kernel*

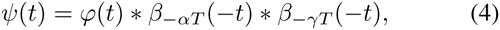

*where β_−αT_ and β_−γT_ are first order E-splines and T is the sampling period.*

Proof.

From the initial filtering operation we obtain

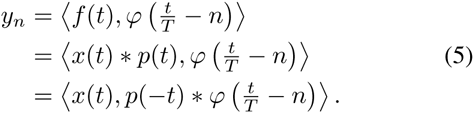

We take finite differences

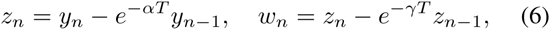

so that we have

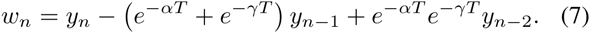

We write *w_n_ =* 〈*x*(*t*)*,h*(*t*)〉 where, from (5) and the linearity of the inner product, we have

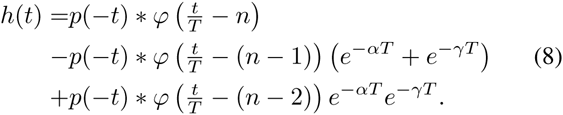

Due to Parseval’s relation *w_n_* can be expressed as

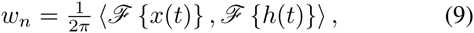

where 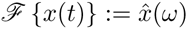 denotes the Fourier Transform (FT) of *x*(*t*). By the linearity of the FT and the FT convolution theorem, we have

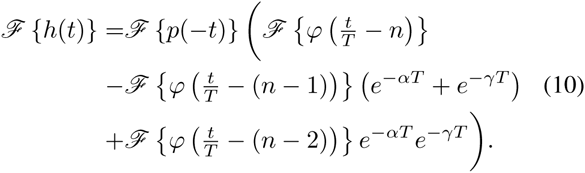

Noting that

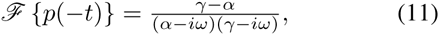

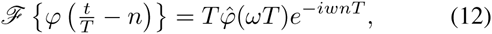

Equation (10) becomes

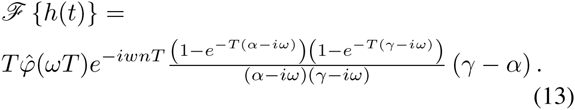

The FT of a time-reversed and scaled E-spline with parameter −*αT* is

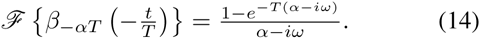

From (12) and (14) it follows that

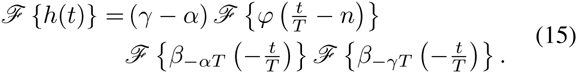

Using Parseval’s relation once more we can write

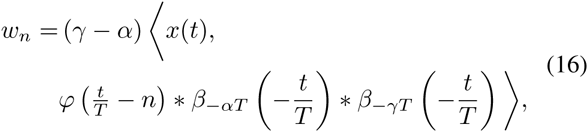

which, with *ψ*(*t*) = *φ*(*t*) * *β*_−_*_αT_*(−*t*) * *β*_−_*_γ__t_*(−*t*), is the statement of the proposition.

### 2.1. Detection of calcium transients in the noisy scenario

The process of detecting the time points of calcium transients from a noisy fluorescence signal is summarised in Algorithm 1 and described in further detail in this section. We model the samples 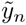 as being corrupted by additive white noise so that we have

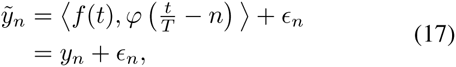
where *ϵ_n_* are i.i.d Gaussian with zero mean and standard deviation *σ*. The sampling kernel *φ* is chosen to be an exponential reproducing kernel. This is defined such that, when it is combined in a weighted sum with shifted versions of itself, it reproduces exponentials:

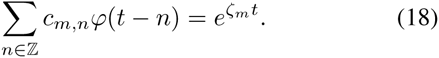

Here, *c_m,n_* are referred to as the coefficients of the kernel. The weighted finite differences in (3) result in noisy samples

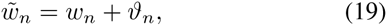
where *ϑ_n_* are Gaussian but no longer i.i.d. We compute sample moments

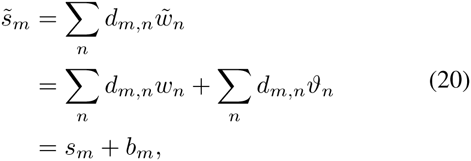
using the coefficients *d_m,n_* of the kernel *ψ* (from (4)) which also reproduces exponentials (see [12]). By choosing *φ* so that *ψ* reproduces exponents *ζ_m_* in the form *ζ_m_* = *ζ*_0_ + *m*λ, we can write *s_m_* in power-sum series form:

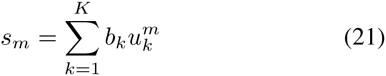
where 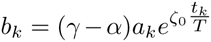 and 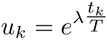. The sample moments *s_m_* are used to construct a Toeplitz matrix 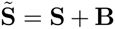, where **S** is the idealised noiseless Toeplitz matrix and **B** is due to the noise *b_m_* corrupting the sample moments. The parameters *u_k_* and thus the time points of the calcium transients *t_k_* are then retrieved from 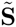 by the matrix pencil method [13]. As the noise has been colored by the filtering process, we first include a pre-whitening step which improves the robustness of the subspace estimation from 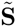.

### 2.2. Pre-whitening to increase robustness to noise

We implement the matrix pencil method to recover the parameters *u_k_*, and thus the time points 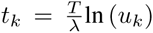, from the noisy Toeplitz matrix 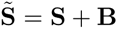. This method performs

#### Algorithm 1: Estimate spike times and amplitudes

**Input***: f(t), K, α, γ*

**Output**: 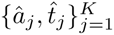

1. Filter: 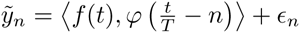
2. Weighted finite differences: 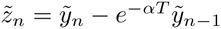
3. Weighted finite differences: 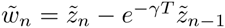
4. Compute sample moments: 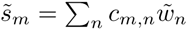
5. Create Toeplitz matrix 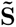 from 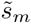
6. Pre-whiten Toeplitz matrix: 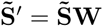
7. Use Matrix Pencil Method to estimate 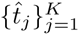 from 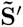
8. Estimate 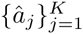 by least squares from samples and resynthesized sample estimates

well when the matrix **B** is white, i.e. **R***_B_* := E [**B***^h^***B**] = a**I** for some real *a*.

The weighted finite differences in Algorithm 1 result in colored noise, such that **R***_B_* ≠ a**I**. We therefore follow the method of Urigüen et al. [14] to pre-whiten 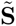. We do this by post-multiplying 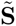 with the matrix 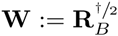 (where ^†/2^ denotes the square root of the pseudo-inverse) to obtain

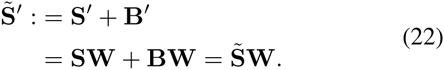

We thus have a noisy Toeplitz matrix 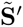 which is corrupted by white noise. The matrix pencil method can then be applied to 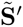 to obtain the same signal parameters *u_k_* as would be obtained from 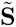 (for a detailed explanation see [15]) whilst maintaining robustness to noise.

### 2.3. Least squares model-order estimation

We implement a least squares model-order estimation framework similar to that proposed by Doğan et al. [17] to estimate the number of spikes in a window of the trace. The training error between the samples and the resynthesized sample estimates is computed for each possible model order *k*. We estimate the model order 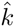 as the value of *k* which minimises the training error, we then estimate the corresponding spike times, according to Algorithm 1. This procedure is completed sequentially in a sliding window along the fluorescence signal. Finally, spike time estimates which are not consistently detected across windows are deemed to be spurious spikes due to noise and are pruned.

## 3. RESULTS

We now demonstrate the performance of the modified FRI algorithm on real GCaMP6s imaging data [16]. The dataset, which is recorded from *in vivo* V1 neurons, contains simultaneous calcium imaging (sampled at 60Hz) and electrophysiology. This allows us to compare estimated spike positions from the calcium imaging data with the ground truth from the electrophysiology. We assess algorithm performance on 3 traces of combined length 678s containing 532 spikes.

A spike is deemed detected if one is estimated within 2 sample widths (0.034s) of the original spike. As the rise of a GCaMP6s transient lasts approximately 15 sample widths, this is a strict target. Of the alternative spike detection algorithms that do compare their spike time estimates against the ground truth, they either tend to not declare the length of their spike detection window [6, 18, 19] or allow a generous time period [20], thereby relaxing the performance metric.

In Figure 1a we show the ROC curves for the modified and original FRI algorithms on one dataset. The true positive rate is the percentage of correctly detected spikes and the false positive rate is the percentage of detected spikes which are not within 2 sample widths of a true spike. It can be seen that the modified FRI algorithm correctly detects a higher proportion of spikes than the original FRI algorithm for all false positive rates in the range of practical interest.

**Fig. 1:**
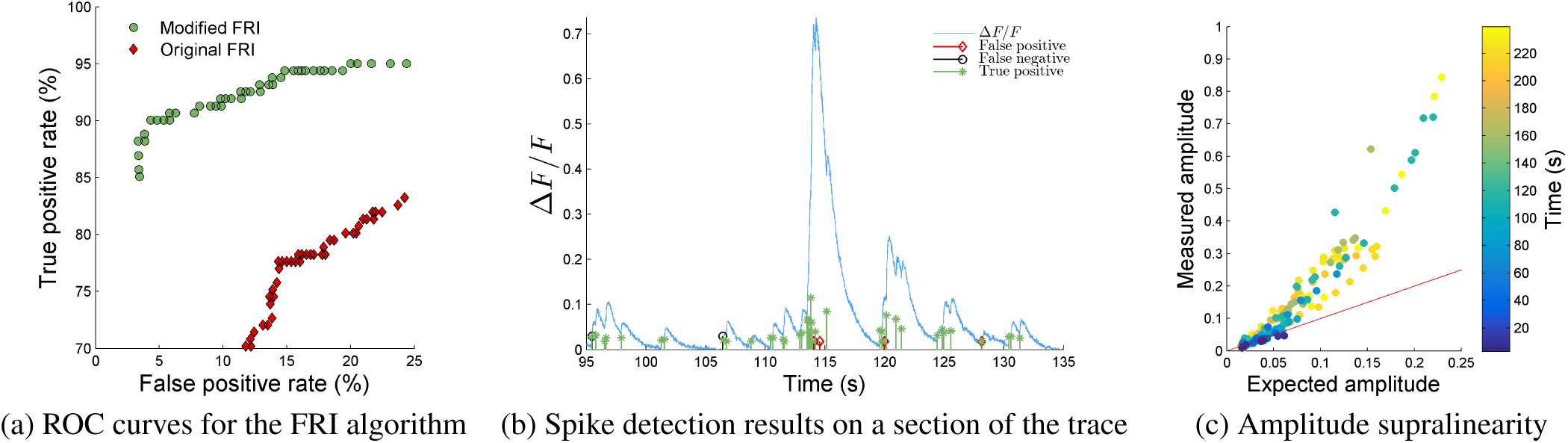
The modified FRI algorithm performs well on a real GCaMP6s dataset [16] of length 239s containing 181 spikes. a) Comparison of the ROC curves of the modified and original FRI algorithms. b) Spike detection performance on a section of the trace. c) The measured amplitude of the fluorescence signal after a spike against the expected amplitude according to a model of uniform fluorescence change per spike. It can be seen that amplitudes sum supralinearly.

Figure 2 displays the average ROC curves over all three datasets for the modified and original FRI algorithms and Vogelstein et al.’s fast non-negative deconvolution algorithm [6], which is commonly used due to its speed and ease of implementation. To compute the average ROC curves a least squares spline fit was computed for each curve and the values of those splines were averaged per algorithm.

**Fig. 2:**
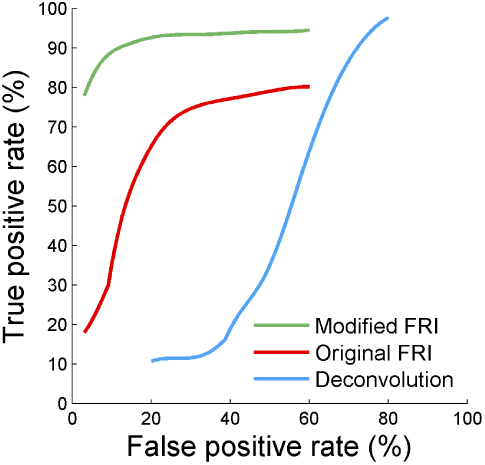
ROC analysis for two variants of the FRI algorithm and the Vogelstein et al. [6] algorithm, averaged over three datasets for which electrophysiological ground truth is available [16]. A spike is deemed detected if one is estimated within 0.034s (2 sample widths) of the true spike position.

Vogelstein et al.’s method detects a high proportion of false positives on this dataset. This is likely due to their assumption of a uniform fluorescence change per spike which, as is shown in Figure 1c and noted in [1], is not true for GCaMP6s. Although the deconvolution algorithm has previously shown good performance on *in vitro* data generated by the synthetic dye OGB-1, it doesn’t perform as well on GCaMP6s data. This demonstrates the necessity of tailoring algorithms to the calcium indicator when performing calcium transient detection. Furthermore, in Figure 2 we can see that the modified FRI algorithm detects a higher proportion of spikes than the original algorithm for all false positive rates.

## 4. CONCLUSION

We extended the FRI framework for spike detection from calcium imaging data to encompass calcium transients with a slow rise, such as those generated by the genetically encoded calcium indicator GCaMP6s. We introduced a noise reduction technique (pre-whitening) and least squares model-order estimation to improve the robustness of the algorithm. On real GCaMP6s data we showed that these modifications improved the spike detection rate of the algorithm compared to the original for all false positive rates in the range of practical interest. Furthermore, on real data we achieve spike detection rates of 90% of electrophysiologically-validated spikes within 2 sample widths (0.034s) of the real spike.

## REFERENCES

[1] T.-W. Chen, T. J. Wardill, Y. Sun, S. R. Pulver, S. L. Renninger, A. Baohan, E. R. Schreiter, R. A. Kerr, M. B. Orger, V. Jayaraman et al., “Ultrasensitive fluorescent proteins for imaging neuronal activity,” Nature, vol. 499, no. 7458, pp. 295–300, 2013.

[2] D. Huber, D. Gutnisky, S. Peron, D. O’Connor, J. Wiegert, L. Tian, T. Oertner, L. Looger, and K. Svoboda, “Multiple dynamic representations in the motor cortex during sensorimotor learning,” Nature, vol. 484, no. 7395, pp. 473–478, 2012.

[3] J. N. D. Kerr, C. P. J. de Kock, D. S. Greenberg, R. M. Bruno, B. Sakmann, and F. Helmchen, “Spatial organization of neuronal population responses in layer 2/3 of rat barrel cortex,” Journal of Neuroscience, vol. 27, no. 48, pp. 13 316–13 328, 2007.

[4] S. R. Schultz, K. Kitamura, A. Post-Uiterweer, J. Krupic, and M. Häusser, “Spatial pattern coding of sensory information by climbing fiber-evoked calcium signals in networks of neighboring cerebellar purkinje cells,” Journal of Neuroscience, vol. 29, pp. 8005–8015, 2009.

[5] B. F. Grewe, D. Langer, H. Kasper, B. M. Kampa, and F. Helmchen, “High-speed in vivo calcium imaging reveals neuronal network activity with near-millisecond precision.” Nature methods, vol. 7, no. 5, pp. 399–405, May 2010.

[6] J. T. Vogelstein, A. M. Packer, T. A. Machado, T. Sippy, B. Babadi, R. Yuste, and L. Paninski, “Fast nonnegative deconvolution for spike train inference from population calcium imaging,” Journal of Neurophysiology, vol. 104, no. 6, pp. 3691–3704, 2010.

[7] J. Oñativia, S. R. Schultz, and P. L. Dragotti, “A finite rate of innovation algorithm for fast and accurate spike detection from two-photon calcium imaging,” Journal of Neural Engineering, vol. 10, no. 4, p. 046017, 2013.

[8] M. Vetterli, P. Marziliano, and T. Blu, “Sampling signals with finite rate of innovation,” Signal Processing, IEEE Transactions on, vol. 50, no. 6, pp. 1417–1428, Jun 2002.

[9] S. P. Peron, J. Freeman, V. Iyer, C. Guo, and K. Svoboda, “A cellular resolution map of barrel cortex activity during tactile behavior,” Neuron, vol. 86, no. 3, pp. 783–799, 2015.

[10] S. Reynolds, J. Oñativia, C. S. Copeland, S. R. Schultz, and P. L. Dragotti, “Spike detection using FRI methods and protein calcium sensors: performance analysis and comparisons,” in 11th international conference on Sampling Theory and Applications (SampTA 2015), Washington, DC, USA, May 2015.

[11] T. Blu, P. L. Dragotti, M. Vetterli, P. Marziliano, and L. Coulot, “Sparse sampling of signal innovations,” Signal Processing Magazine, IEEE, vol. 25, no. 2, pp. 31–40, 2008.

[12] M. Unser and T. Blu, “Cardinal exponential splines: Part I—Theory and filtering algorithms,” IEEE Transactions on Signal Processing, vol. 53, no. 4, pp. 1425–1438, April 2005.

[13] Y. Hua and T. K. Sarkar, “Matrix pencil method for estimating parameters of exponentially damped/undamped sinusoids in noise,” Acoustics, Speech and Signal Processing, IEEE Transactions on, vol. 38, no. 5, pp. 814–824, 1990.

[14] J. A. Urigüen, T. Blu, and P. L. Dragotti, “FRI sampling with arbitrary kernels,” Signal Processing, IEEE Transactions on, vol. 61, no. 21, pp. 5310–5323, 2013.

[15] J. A. Urigüen, “Exact and approximate strang-fix conditions to reconstruct signals with finite rate of innovation from samples taken with arbitrary kernels,” Ph.D. dissertation, Imperial College London, 2013. [Online]. Available: http://hdl.handle.net/10044/1Z12817

[16] GENIE project, Janelia Farm Campus, HHMI; Karel Svoboda (contact). (2015). Simultaneous imaging and loose-seal cell-attached electrical recordings from neurons expressing a variety of genetically encoded calcium indicators. CRCNS.org. http://dx.doi.org/10.6080/K02R3PMN.

[17] Z. Doğan, C. Gilliam, T. Blu, and D. Van De Ville, “Reconstruction of finite rate of innovation signals with model-fitting approach,” Signal Processing, IEEE Transactions on, vol. PP, no. 99, pp. 1–1, 2015.

[18] F. D. Andilla and F. A. Hamprecht, “Learning multilevel sparse representations,” in Advances in Neural Information Processing Systems, 2013, pp. 818–826.

[19] F. D. Andilla and F. A. Hamprecht, “Sparse space-time deconvolution for calcium im*age analysis*,” in Advances in Neural Information Processing Systems, 2014, pp. 64–72.

[20] H. Lütcke, F. Gerhard, F. Zenke, W. Gerstner, and F. Helmchen, “Inference of neuronal network spike dynamics and topology from calcium imaging data,” Frontiers in neural circuits, vol. 7, 2013.

